# The phers R package: using phenotype risk scores based on electronic health records to study Mendelian disease and rare genetic variants

**DOI:** 10.1101/2022.06.07.495133

**Authors:** Layla Aref, Lisa Bastarache, Jacob J. Hughey

**Affiliations:** Department of Biomedical Informatics, Vanderbilt University Medical Center, Nashville, Tennessee, USA; Program in Chemical and Physical Biology, Vanderbilt University School of Medicine, Nashville, Tennessee, USA

## Abstract

Electronic health record (EHR) data linked to DNA biobanks are a valuable resource for understanding the phenotypic effects of human genetic variation. We previously developed the phenotype risk score (PheRS) as an approach to quantify the extent to which a patient’s clinical features resemble a given Mendelian disease. Using PheRS, we have uncovered novel associations between Mendelian disease-like phenotypes and rare genetic variants, and identified patients who may have undiagnosed Mendelian disease. Although the PheRS approach is conceptually simple, it involves multiple mapping steps and was previously only available as custom scripts, limiting the approach’s usability. Thus, we developed the phers R package, a complete and user-friendly set of functions and maps for performing a PheRS-based analysis on linked clinical and genetic data. The package includes up-to-date maps between EHR-based phenotypes (i.e., ICD codes and phecodes), human phenotype ontology (HPO) terms, and Mendelian diseases. Starting with occurrences of ICD codes, the package enables the user to calculate phenotype risk scores, validate the scores using case-control analyses, and perform genetic association analyses. By increasing PheRS’s transparency and usability, the phers R package will help improve our understanding of the relationships between rare genetic variants and clinically meaningful human phenotypes.

**Availability:** The phers R package is free and open-source, and available on CRAN and at https://phers.hugheylab.org.

**Supplementary information:** Supplementary data are available at *Bioinformatics* online.

## 1 Introduction

Rare diseases affect an estimated 300-400 million people worldwide (Nguengang Wakap *et al*., 2020). This large disease burden is distributed among an estimated 5,000-10,000 rare diseases, of which 70-80% may be genetic in origin, most of these monogenic/Mendelian (Amberger *et al*., 2015). The wide variability in these estimates gives an idea of how challenging such rare genetic diseases are to study (Haendel *et al*., 2020).

Much of our knowledge of Mendelian diseases derives from labor-intensive, family-based studies of meticulously phenotyped individuals, which have revealed that these diseases produce an array of symptoms and are often caused by rare genetic variants having large effect sizes. However, genome-wide association studies (GWASs) have typically examined phenotypes independently, and until recently, focused on common variants having modest effect sizes. As a result, the phenotypic effects of many rare variants, even those in Mendelian genes, remain uncertain.

The growth of DNA biobanks linked to electronic health records (EHRs) has created an exciting opportunity to study genotype-phenotype relationships relevant to human health. To enable high-throughput, EHR-based studies of Mendelian disease, we previously developed an approach that aggregates an individual’s clinical diagnoses based on known features of a given Mendelian disease and summarizes the evidence as a phenotype risk score (PheRS) (Bastarache *et al*., 2018, 2019). Using PheRS, we have discovered and subsequently validated novel associations between Mendelian disease-like phenotypes and rare variants in Mendelian genes. Our findings have also identified patients whose clinical diagnoses may have a genetic cause, blurring the line between Mendelian and complex disease. Although PheRS is conceptually simple, it involves multiple mapping steps between various ontologies, e.g., between EHR-based phenotypes and Mendelian diseases. Furthermore, previous implementations were only available as custom scripts and database queries specific to our institution’s data and our analyses. To address this lack of usability, we developed the phers R package, a user-friendly and open-source implementation of the PheRS approach. Here we describe each component of the package and show a case study of using the package to perform a PheRS-based analysis of linked clinical and genetic data.

## 2 Methods

The main inputs for a typical PheRS-based analysis are demographics, occurrences of International Classification of Disease (ICD) codes, and genotypes for each person in the cohort. The typical steps are described below.

### 2.1 Mapping ICD codes to phecodes

The phenotype risk score relies on phecodes (Denny *et al*., 2013), custom groupings of ICD codes that reduce granularity and unify ICD-9 and ICD-10 codes. The package uses phecode version 1.2 to map ICD occurrences to phecode occurrences. Users can exclude from the mapping any ICD codes that indicate genetic diagnosis of Mendelian disease, a useful feature when comparing scores of diagnosed and undiagnosed individuals.

### 2.2 Assigning weights to phecodes

By default, the package assigns a weight to phecode *j* of 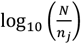, where *N* is the total number of individuals in the cohort and *n*_*j*_ is the number of individuals with at least one occurrence of phecode *j*. The package also includes pre-calculated weights based on EHR data from patients with at least one visit at Vanderbilt University Medical Center (∼3 million individuals; 53% females; range of age at first visit: 0-90 years), which may be useful when the cohort provided by the user is small and/or its phecode prevalences do not reflect those in the population of interest.

### 2.3 Defining disease patterns as groups of phecodes

By default, the package groups phecodes with Mendelian diseases based on two maps: the first describing the clinical features of each Mendelian disease in terms of human phenotype ontology (HPO) terms (Köhler *et al*., 2021) and the second linking HPO terms to phecodes. Alternatively, users can provide a custom grouping of phecodes with diseases.

For the first map, we used HPO annotations for diseases in the Online Mendelian Inheritance in Man (OMIM) catalog. We downloaded the map of OMIM disease IDs and HPO terms (https://hpo.jax.org/app/download/annotation) and the human gene map (morbidmap.txt, https://omim.org/downloads). Using these resources, we created a table linking genes, inheritance patterns, and phenotypic features of Mendelian diseases. We excluded disease-gene pairs having a multifactorial, somatic, or unspecified inheritance pattern, as well as pairs marked as provisional. For the second map, we manually linked HPO terms and phecodes, expanding the map from our prior work (Bastarache *et al*., 2019) to cover all Mendelian diseases included in the first map described above. Of the 6,451 unique HPO terms linked to an OMIM disease ID, we matched 4,189 to a phecode. We deemed only 278 of these matches exact, with the majority having a broader phecode linked to a more specific HPO term. Researchers should consider this expanded map a starting point, and where possible validate it on their own data.

After merging these two maps, we further filtered the result to only include Mendelian diseases having at least three unique phecodes. The final phecode-to-disease grouping in the package includes 4,305 unique Mendelian diseases and 984 unique phecodes.

### 2.4 Calculating the phenotype risk score

Given a set of weights, phecode occurrences, and grouping of phecodes with diseases, the package calculates the phenotype risk score *s*_*i,k*_ for person *i* and disease *k* as

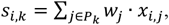

where *P*_*k*_ is the set of phecodes for disease *k, w*_*j*_ is the weight for phecode *j*, and *x*_*i,j*_ is 1 if person *i* has an occurrence of phecode *j* and 0 otherwise. The package can also calculate residual phenotype risk scores based on a linear model that can include terms for demographics and other individual-level variables. The residual scores are standardized to have unit variance, and are thus more comparable from one disease to another.

### 2.5 Identifying individuals diagnosed with genetic disease

The package can help validate that individuals diagnosed with a given Mendelian disease have a higher phenotype risk score for that disease compared to undiagnosed individuals. Such validation is an important step to perform prior to testing new hypotheses.

To identify diagnosed (cases) and undiagnosed individuals (controls) for each Mendelian disease, the package uses ICD occurrences provided by the user and a map (included in the package) of diagnostic ICD codes and Mendelian diseases. By default, individuals having a diagnostic ICD code on at least two distinct dates are defined as cases, on zero dates as controls, and on one date as neither.

To create the map, we used the Orphanet Rare Disease Ontology (ORDO) (Jupp *et al*., 2015), extracting ICD-10 codes and OMIM disease IDs from the hasDbXref column. We manually reviewed these pairs and then mapped ICD-10 to ICD-9 codes using the crosswalk. We kept in the map diseases having both ICD-9 and ICD-10 codes. The final map includes only 27 Mendelian diseases, as most lack a specific ICD code.

### 2.6 Running genetic association studies

Given scores and genotypes, the package can perform genetic association tests based on linear regression and either an additive, dominant, recessive, or genotypic model. The model can include terms for demographics and other individual-level variables. The user must specify which genetic variants should be tested for which Mendelian diseases. For each disease-variant pair, the package reports the counts of individuals having each genotype and the model’s summary statistics.

## 3 Results

To demonstrate the utility of the phers R package, we used it to analyze de-identified EHR and genetic data from Vanderbilt University Medical Center (VUMC). We first assembled a cohort of 3,054,539 individuals having at least one visit at the medical center. We defined cases and controls for 27 Mendelian diseases based on ICD-9 and ICD-10 code occurrences (using the default setting of cases having a diagnostic ICD code on at least two distinct dates and controls having none). After mapping ICD code occurrences to phecodes, we calculated each individual’s phenotype risk score for each disease (Table S1). We also calculated residual scores based on a linear model having terms for biological sex, age at first visit, and amount of time between first and last visits (Fig. 1A). For the 20 Mendelian diseases that had at least 50 cases, the corresponding phenotype risk scores tended to be higher in cases than in controls (p < 1.4.10^−66^ based on linear regression adjusted for biological sex, age at first visit, and amount of time between first and last visits). Although the presence of diagnostic ICD codes does not always indicate a definitive genetic diagnosis, these results suggest that the scores calculated by the phers package on our data do reflect the risk of Mendelian disease.

**Fig. 1.**
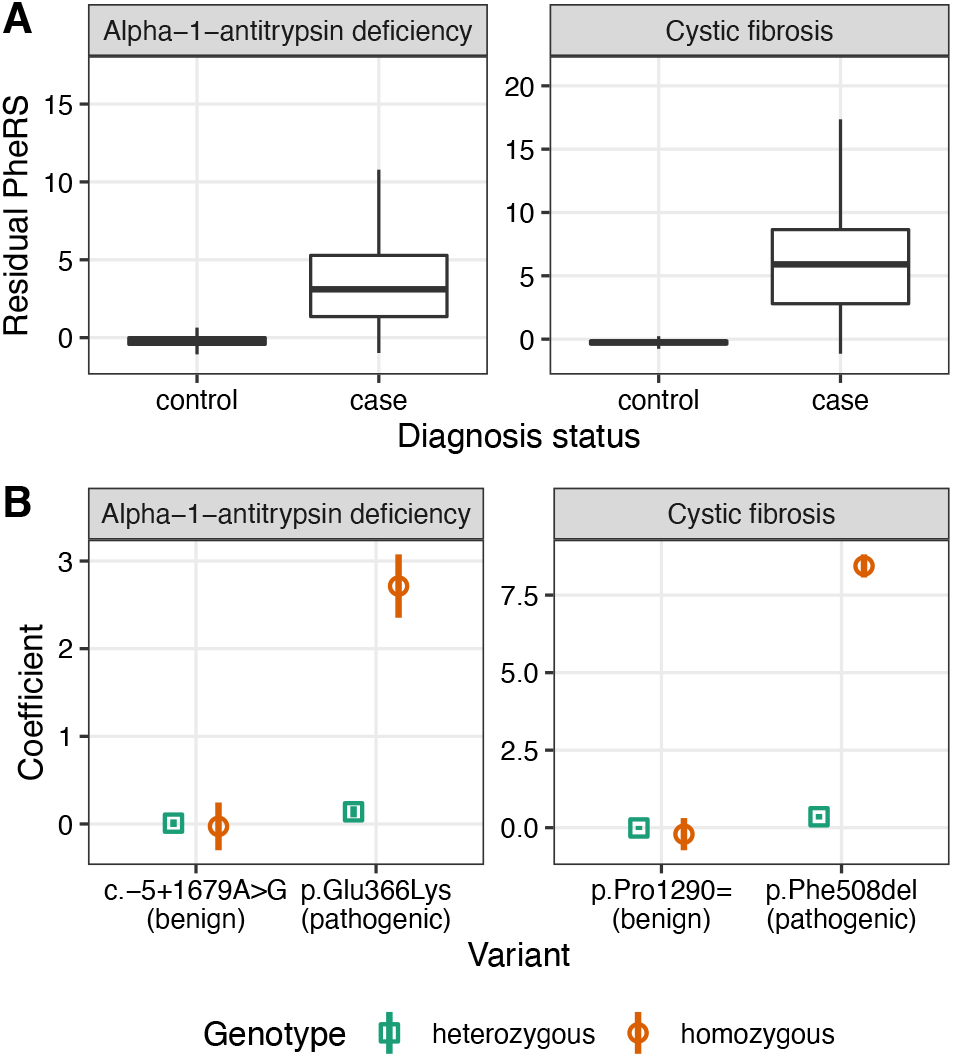
Example PheRS-based analysis. **(A)** Boxplots of residual phenotype risk scores for cases and controls of alpha-1-antitrypsin deficiency (612 cases and 3,053,472 controls) and cystic fibrosis (1,374 cases and 3,052,073 controls). For clarity, outliers are not shown. (B) Coefficients (*points*) and corresponding 95% confidence intervals (*vertical lines*) of associations between phenotype risk scores for the given disease and selected genetic variants. Annotations (*benign* or *pathogenic*) are based on ClinVar. Numbers of heterozygous and homozygous individuals for each variant are in Table S2.

We next sought to use the package to quantify associations between phenotype risk scores and rare variants in Mendelian genes. We assembled a cohort of 69,416 adults of European ancestry genotyped on the MultiEthnic Genotyping Array (MEGA) and having linked EHR data at VUMC. For each of two Mendelian diseases (cystic fibrosis and alpha-1antitrypsin deficiency), we selected a pathogenic variant and a benign variant based on ClinVar (Landrum *et al*., 2020). For each disease-variant pair, we fit a genotypic model having additional terms for biological sex, age at first visit, and amount of time between first and last visits. As expected for a recessive inheritance pattern, the coefficients of heterozygosity and homozygosity for the benign variants, as well as the coefficients of heterozygosity for the pathogenic variants, were near zero. In contrast, the coefficients of homozygosity for the pathogenic variants were strongly positive (Fig. 1B and Table S2). These results illustrate how the scores calculated by the package can reveal the phenotypic effects of rare genetic variants.

## 4 Discussion

The PheRS approach and the phers package do have limitations. First, as with any approach based on observational data, care is warranted when interpreting statistical associations, as they may be due to bias and not causality. Second, the performance of the various maps, particularly the newly expanded HPO-to-phecode map, may vary depending on local ICD billing practices. Third, despite the importance of validating phenotypes via case-control analysis, cases for the vast majority of Mendelian diseases cannot be identified based solely on ICD codes, which may necessitate use of other data types or manual chart review.

Overall, however, the phers package provides a solid platform for large-scale, high-throughput studies of Mendelian disease that can complement detailed family-based studies. As the amount of linked EHR and genetic data grows, the phers package can help researchers use these data to gain knowledge about rare diseases and their underlying genetics.

## Acknowledgements

We thank Josh Schoenbachler and Elliot Outland for feedback on the code for the package and the analysis.

## Data availability

Access to individual-level EHR and genotype data is IRB-restricted. Code and summary-level results are at https://doi.org/10.6084/m9.figshare.20016728.

